# PolyCRACKER, a robust method for the unsupervised partitioning of polyploid subgenomes by signatures of repetitive DNA evolution

**DOI:** 10.1101/484832

**Authors:** Sean P. Gordon, Joshua J. Levy, John P. Vogel

**Affiliations:** DOE Joint Genome Institute, 2800 Mitchell Dr., Walnut Creek, CA 94598; University of California Berkeley, Berkeley, CA

**Author notes:** contributed equally to this work (Sean P. Gordon, Joshua J. Levy).

**Keywords:** allopolyploid, k-mer, transposon, binning, evolution, repetitive DNA, subgenome, wheat, tobacco, metagenome

## Abstract

Existing methods for assigning sequences to individual species from pooled DNA samples rely on differences in genome properties like GC content or sequences from related species. These approaches do not work for closely related species where gross features are indistinguishable and related genomes are lacking. We describe a method and associated software package that uses rapidly evolving repetitive DNA to circumvent these limitations. By using short, repetitive, DNA sequences as species-specific signals we separated closely related genomes without any prior knowledge. This approach is ideal for separating the subgenomes of polyploid species with unsequenced or unknown progenitor genomes.

## Introduction

Researchers studying two traditionally distinct areas of biology, metagenomics and polyploid genome evolution, face a similar technical challenge: How do you separate closely related genomes or subgenomes from a single sample? Current approaches consist of either supervised binning, requiring a database of known genomes, or unsupervised binning using general genome characteristics as a ‘genome signature’ to identify and separate DNA sequences from different species/subgenomes [1, 2]. Genome signatures can be any kind of bias (e.g. similarity to known sequences, short nucleic acid sequences, GC content, sequence depth) that differs between species or subgenomes. In particular, di- and tetra-nucleotide frequencies along with contig co-abundance (a given species may have a distinct abundance relative to others in a metagenome sample) can be used to successfully bin sequences within metagenomes [3]. These coarse approaches work best with highly divergent taxa and there is much room for improvement in sensitivity and accuracy when grouping sequences appropriately at the species level. Furthermore, subgenomes within allopolyploids have the same sequence depth profile and similar di- and tetra-nucleotide frequencies, making these features uninformative. In fact, orthologous gene sequences between the subgenomes in an allopolyploid (homeologous sequences) are far more similar to each other than they are to other sequences within the same subgenome.

Polyploidization has played a large role in the evolution of all flowering plants and many extant species are recent polyploids [4]. In addition, polyploidization has played a role in the evolution of many fungal, fish and amphibian species [5, 6]. A large fraction of polyploid species are derived from the hybridization of different species and are termed allopolyploids. Understanding the evolution and regulation of polyploid genomes requires the creation of accurate assemblies for each subgenome. The high level of sequence similarity between subgenomes makes this particularly challenging and even state-of-the-art genome assembly algorithms often collapse and/or incorrectly interweave chunks of homeologous chromosomes. Significantly, without high resolution genetic maps or related data these errors remain undetected. These errors are not necessarily resolved by long-read technologies, since such assemblies are still fragmented in many thousands of pieces [7]. Lack of subgenome resolution for allopolyploid genomes is a major obstacle to studying the origin, evolution, and functional analysis of allopolyploid species [8].

For some allopolyploid species, extant diploid species similar to the true progenitors can be used to disentangle subgenomes [7]. However, for many allopolyploids [9, 10], the diploid parents are unknown or extinct. Even when extant progenitor-like species exist, they may not well reflect the genomes of the individuals that gave rise to the allopolyploid. Thus, unguided methods for identification and extraction of subgenomes within allopolyploids would be extremely useful. Recently a phasing method was developed for the highly heterozygous sweet potato genome [11]. However, this method is limited to highly polymorphic genomes and is restricted to single-copy sequence present in each subgenome that can be accurately genotyped.

Deliberate integration of synthetic transposons with molecular barcodes is a common experimental approach to label subpopulations of DNA or cells to allow subsequent retrieval from complex mixtures [12]. Nature’s equivalents of synthetic molecular barcodes include transposons, viruses, and other classes of repetitive DNA. These elements rapidly proliferate and evolve in natural populations, thus labeling the recent evolutionary history of an organism’s genome (Fig. 1) [13]. Transposon families may dramatically expand and contract over short periods of evolutionary time, during which they may also significantly change in sequence identity. For example, the size of the *Oryza australiensis* genome doubled in just a few million years due to the amplification of a few LTR retrotransposon families [14]. Removal of LTR retrotransposons by illegitimate recombination can eliminate megabases of sequence over similar timescales [13, 15]. Thus, two recently diverged species may quantitatively differ in the frequency of transposon families they harbor and the sequence identity of those families.

**Figure 1.**
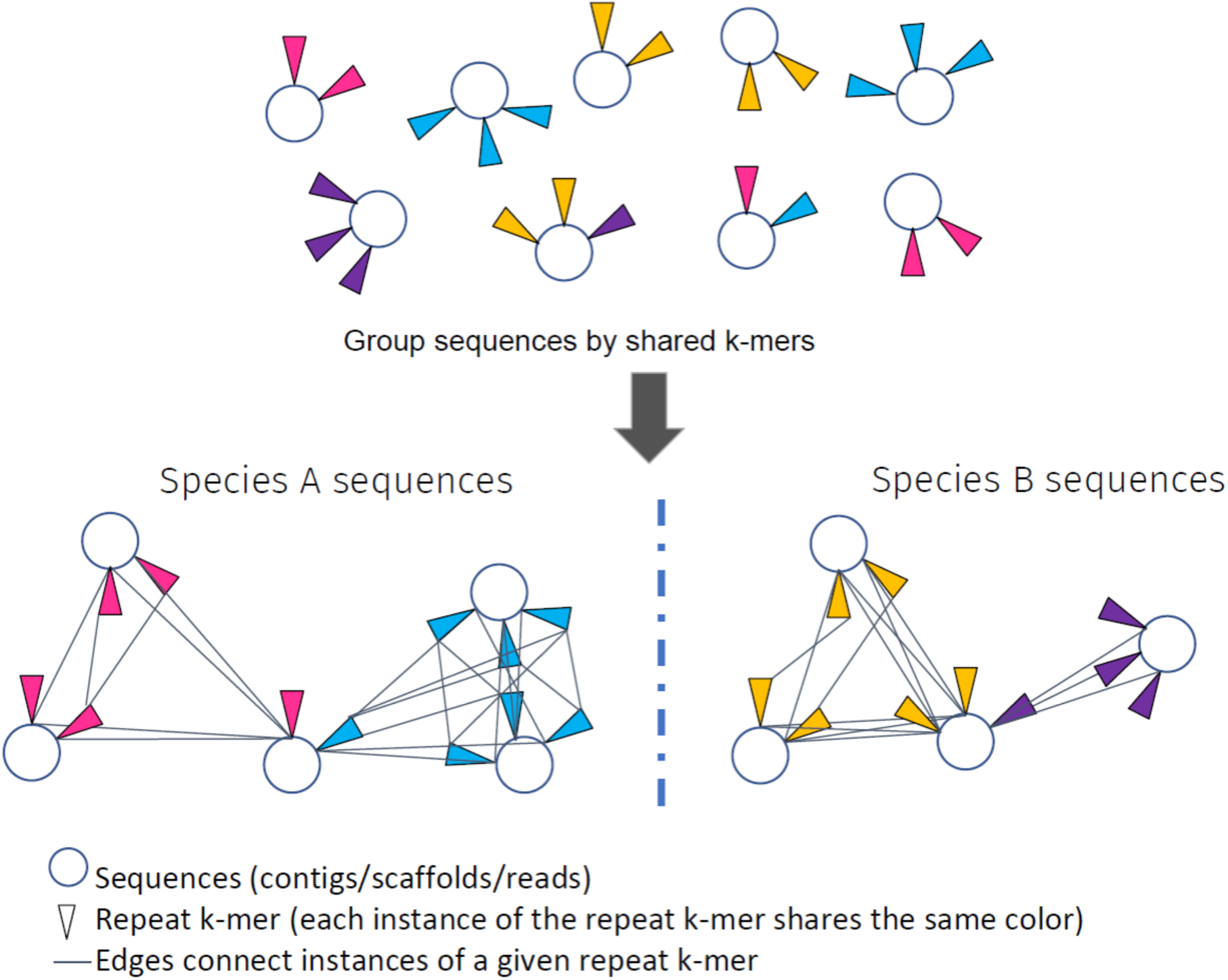
Overview of the strategy for determining species of origin for sequences based on their profile of repetitive k-mers. The algorithm takes sequences from a sample containing one or more species. Each species has both common and divergent repetitive k-mers scattered throughout its genome. Each k-mer is indicated by a different color (for clarity, only divergent k-mers are shown). We construct a graph where sequences are nodes and shared instances of repetitive k-mers are edges. By connecting all instances of a given repetitive k-mer we are able to connect and subsequently group sequences derived from a common species.

In the context of allopolyploids, repeats in the parental species change and proliferate independently after those species diverge creating species-specific genomic signatures. Once reunited within a single cell of an allopolyploid, these repeats are free to proliferate between the subgenomes and these additional copies are specific to the allopolyploid but not the progenitor species. If additional subgenomes are added via subsequent additional hybridizations, then the initial allopolyploid-specific repeat signatures may distinguish initial subgenomes from subgenomes added during later rounds of inter-specific hybridization. Fortuitously, the process of interspecific hybridization may in fact trigger virus and transposon proliferation [16]. Thus, naturally occurring repetitive DNA can be used as molecular barcodes to distinguish genomes and subgenomes.

Here we describe an algorithm that identifies natural molecular barcodes that act as species of origin labels for the DNA sequences in which they reside. Our method creates and partitions a graph in which sequences are connected by edges, corresponding to specific signatures of repetitive DNA inferred from 11-131 nucleotide sequences (k-mers). Our method allows accurate separation of the subgenomes of polyploid species without prior knowledge of the genomes or a database of known sequences. We further built a toolkit of functions, implemented as a python package called polyCRACKER (polyploid Cluster Repeats by Ancestral Common K-mer Estimation and Retrieval), for the automatic binning of sequence scaffolds from polyploid genomes. Our method draws its inspiration from other unsupervised algorithms for binning sequences commonly applied to metagenome datasets, but by focusing on repetitive k-mers with faster rates of divergence between two species, rather than all k-mers or a random sampling of k-mers, our method is not confounded by the overall high sequence similarity between homeologous sequences in allopolyploids or sequencing datasets containing genomes from multiple closely related species.

## Results

We designed an efficient unsupervised learning algorithm to automatically identify and extract DNA sequences within complex mixtures based on the shared presence of short repeated DNA sequences (k-mers) that act as naturally occurring molecular barcodes (Fig. 1). A sample containing a mixture of species is sequenced and assembled to scaffolds that are unordered with respect to their species of origin. Most scaffolds contain multiple k-mers derived from repetitive DNA elements. We construct a graph where sequences are nodes and their profile of shared repeat-k-mers are edges. By connecting all instances of a given repetitive k-mer, we are able to connect and subsequently group sequences derived from a common species (Fig. 1). It is notable that while both high and low copy k-mers may be specific to one species versus another, only high copy k-mers provide the multiple edges between multiple different sequence nodes necessary to group sequences together by species of origin.

### polyCRACKER method for unsupervised partitioning of sequence into species bins

Unsupervised partitioning of sequences proceeds in two phases: 1) initial partitioning of sequences into species bins via a nearest neighbor’s graph of repeat-k-mers and 2) signal amplification from groups identified in the first phase.

Initial partitioning of sequences into species bins can be broken down into three main steps.

a. Repeat-k-mers are selected based on the number of times they appear throughout the genome. Typically, fixing the genome size, as the k-mer size is increased, the threshold for the lowest allowable k-mer frequency should be reduced. As the genome size is increased, these thresholds can be relaxed. This balance is based on the requirement that any given subsequence (contig or scaffold) in the genome must have one or more repeat-k-mers associated with it in order for it to be portioned into a subgenome bin.
b. Sequences from a sparse count matrix of genome sequences versus k-mers are projected into a lower dimensional space, where the distance between points/scaffolds is indicative of the degree of shared repetitive k-mer content.
c. A nearest neighbors graph is constructed from these projected data points. Each node represents a sequence, while each edge demonstrates that two sequences share a highly similar distribution of repetitive k-mers. In this sense, an edge is a proxy for connecting the instances of shared repeat-k-mers. The scaffolds are partitioned into bins by cutting this graph at regions with a low density of connections.

The signal amplification phase consists of two iterative steps that recruit previously unclassified scaffolds to the appropriate subgenome/species bins.

d. Differential repeat-k-mers are identified between the initially extracted subgenomes/species bins. The frequencies of those differential repeat-k-mers within all genome sequences is used to recruit additional sequences into respective species/subgenome bins.
e. A new set of differential repeat-k-mers pertaining to the new subgenomes/species bins are identified, and are once again used assign sequences to a species/subgenome. These steps are iterated until the new species/subgenome bins converge towards a fixed size.

### Polyploid genome analysis

The creation of polyCRACKER was motivated by prior sequence binning applications, particularly in the unsupervised metagenomics binning field. However, tools designed for unsupervised binning of metagenomes either require input that is not relevant for allopolyploid analysis, such as multiple samples from different microbial environments, or rely too heavily on large overall differences in k-mer frequency that are typically not found between highly similar allopolyploid subgenomes. However, as discussed above, it has been shown that the profile of repetitive k-mers may vary considerably between allopolyploid subgenomes. We therefore investigated whether a method specifically focusing on the tracking of repeat-k-mers in allopolyploid genomes would enable database-free species-level binning. Initial results on simulated datasets representing mixtures of closely related microbial genomes showed promise, even when these genomes had relatively low levels of repetitive DNA (Supplementary Data, Fig. 1, Tables 1,2). These results encouraged us to turn our attention to the unsupervised separation of subgenomes from larger, more complex allopolyploid genomes.

**Table 1:** Increasing k-mer length decreases the ability of polyCRACKER to assign *N. tabacum* genome sequences to a subgenome. The threshold for minimum counts for k-mers scales inversely to the k-mer length.

| Kmer length | Minimum k-mer count threshold | Cohen's Kappa: unambiguous; all sequences | Amount Sequence Agreement / Total Genome | Number of k-mers used |
| --- | --- | --- | --- | --- |
| 15 | 150 | 0.998, 0.991 | 0.868 | 929,119 |
| 23 | 150 | 0.998, 0.989 | 0.867 | 296,059 |
| 26 | 150 | 0.998, 0.985 | 0.865 | 195,362 |
| 33 | 120 | 0.998, 0.986 | 0.866 | 108,370 |
| 43 | 100 | 0.997, 0.975 | 0.859 | 35,058 |
| 53 | 80 | 0.993, 0.996 | 0.826 | 13,061 |

**Table 2:** Unsupervised separation of subgenomes from polyploid plants using PolyCRACKER’s differential k-mer analysis.

| Species | Sub sequence length | k-mer length | Genome Size | Cohen's Kappa (only unambiguous) | Cohen's Kappa (including ambiguous) | Sequence in agreement / total sequence |
| --- | --- | --- | --- | --- | --- | --- |
| *N. tabacum* (including unanchored sequences) | 100 kb | 26 | 3.8 Gb | 0.999 | 0.984 | 0.992 |
| *N. tabacum* (only anchored sequences) | 250 kb | 26 | 2.9 Gb | 0.998 | 0.985 | 0.865 |
| *N. tabacum* (only anchored sequences), repeat elements | 250 kb | n/a | 2.9 Gb | 0.995 | 0.87 | 0.795 |
| *Triticum aestivum* | 250 kb | 26 | 15.3 Gb | 0.993 | 0.993 | 0.997 |

### Subgenome analysis of the allotetraploid plant, *Nicotiana tabacum*

*Nicotiana tabacum* is thought to have formed less than 200,000 years ago by the hybridization of two diploid species similar or identical to *Nicotiana sylvestris* (source of the S-subgenome) and *N. tomentosiformis* (source of the T-subgenome). The genome of *N. tabacum* is slightly smaller than the combined sizes of its diploid progenitors due to the preferential loss of repetitive sequence from the T-subgenome [17]. Initial studies comparing the *N. tabacum* genome to the genomes of *N. sylvestris* and *N. tomentosiformis* assigned about 80% of the assembly to the S- or T-subgenomes [7, 18]. A recent study used optical and genetic maps to assign 64% of the assembled *N. tabacum* sequence to pseudomolecules [7]. A significant complicating factor in the separation of the S- and T-subgenomes is the large number of translocations that have occurred between the S- and T-progenitor chromosomes [4]. Thus, identifying and separating the ancestral S- and T-subgenomes from the modern allopolyploid requires classification of alternating stretches of contiguous DNA within chromosomes and not simply labeling of whole chromosomes.

We created an initial dataset for testing and verification of polyCRACKER for polypoid analysis by splitting the sequences within the pseudomolecule-anchored portion of the *N. tabacum* genome into non-overlapping 250 kb segments. This later allowed us to visualize subgenome classification with respect to position within pseudomolecules. This was particularly important for *N. tabacum* due to the numerous translocations that have occurred between subgenomes and the uncertainty of whether polyCRACKER could accurately assign progenitor of origin to pseudomolecules containing sequences from different progenitors. We validated the assignment of progenitor of origin by polyCRACKER by using, in this case, the known (and also assembled) progenitors of the allopolyploid, *N. sylvestris* and *N. tomentosiformis.* We quantified agreement between polyCRACKER subgenome assignments and the subgenome assignments determined by comparison to the progenitor genome sequences. Relationships between *N. tabacum* sequences are depicted in PCA plots (Fig. 2a,b) and a spectrally embedded graph (Fig. 2c). Each data point in the PCA plot corresponds to a 250 kb subsequence and the proximity of the points indicates the degree of shared repetitive k-mers. Points were colored by their inferred progenitor of origin via polyCRACKER (Fig. 2a) or by comparison to the diploid progenitor genomes (Fig. 2b). Subsequences were linked to their 20 nearest neighbors to create a network graph. The energy minimization of a non-repulsive force-directed graph of the 20 nearest neighbors yields spectrally embedded data (spectral embedding/laplacian eigenmaps); polyCRACKER groups subsequences by clustering the spectral embedding of the dimensionality reduced data. This is depicted in Fig. 2c, in which nodes (sequences) are colored according to their species of origin assignment based on comparison to the diploid progenitor genomes. Spectral clustering performs k-means clustering on the spectrally embedded data in Fig. 2c, analogous to making cuts in the weakest links in the graph.

**Figure 2.**
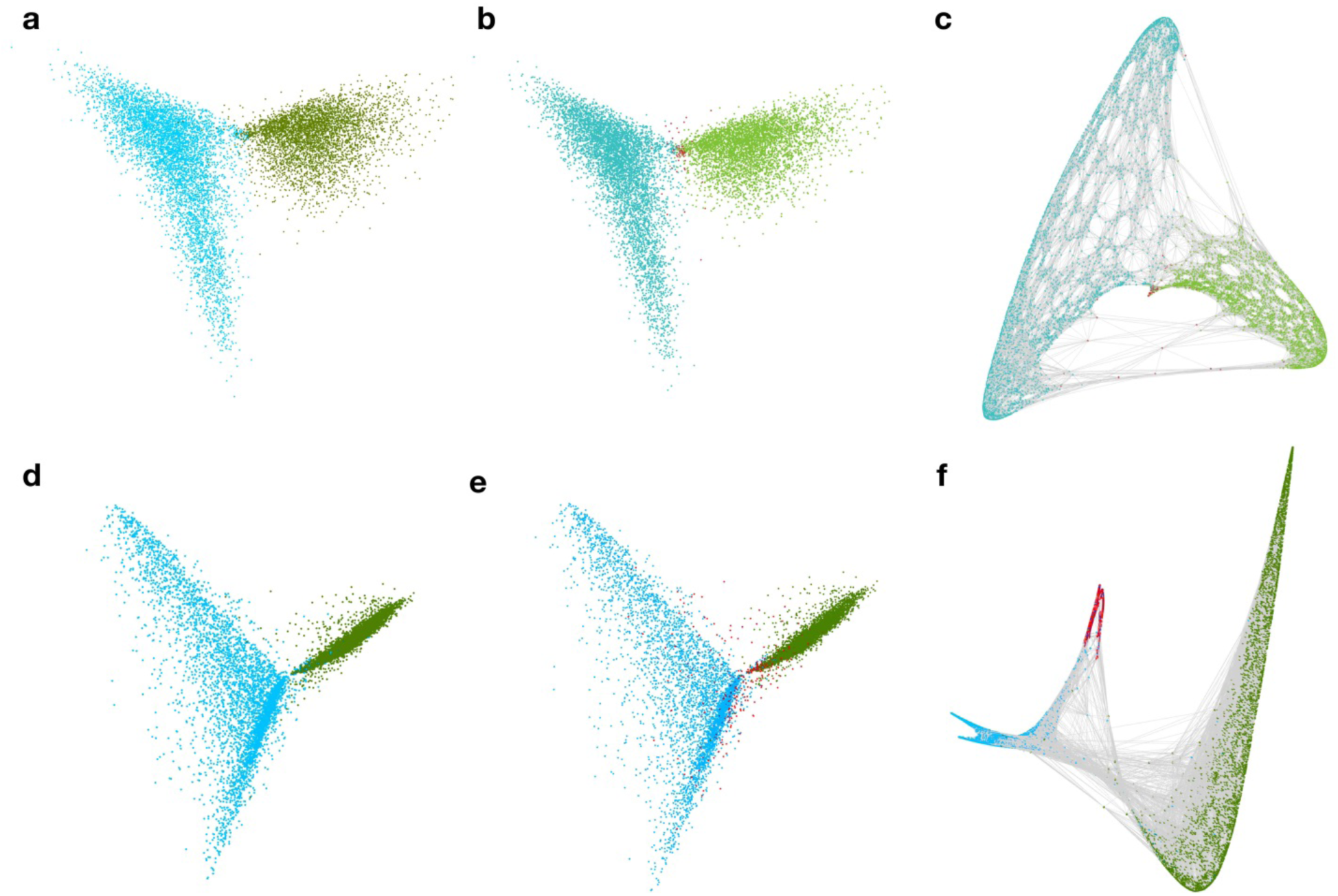
Unsupervised grouping of 250 kb fragments derived from the pseudomolecule-anchored portion of the *N. tabacum* genome by polyCRACKER using repetitive k-mers and intact repeats. (**a** and **b**) PCA of repetitive k-mer count matrix of k-mers versus 250 kb genome segments for *N. tabacum.* S and T genomes are colored green and blue respectively, ambiguous segments are labeled red. (**a**) Sequences labeled by similarity to the known progenitor-like species. (**b**) Sequences labeled by polyCRACKER’s k-mer analysis. Note that **a** and **b** are nearly identical. (**c**) Spectral embedding of *N. tabacum* dimensionality reduced information, in which edges (grey lines) represent shared repetitive k-mer profiles) that connect sequences. Sequences labeled by similarity to the known progenitor-like species. (**d-f**) Analogous unsupervised grouping as above, but using informative and differential repeats rather than k-mers. Color labels as described above. (**d**) Sequences labeled by similarity to the known progenitor-like species. (**e**) Sequences labeled by polyCRACKER’s repeat analysis. Note that **a,b,d** and **e** are nearly identical. (**f**) Spectral embedding of *N. tabacum* dimensionality reduced information, in which edges (grey lines) represent shared repeat element profiles that connect sequences. Sequences labeled by similarity to the known progenitor-like species. Note the increased number of ambiguous subsequences in **f** as compared to c is due to the fact that there are fewer repeats than k-mers in any given scaffold. This is analogous to the effect of increasing k-mer size substantially.

PolyCRACKER correctly classified 99.3 percent of sequence estimated to belong to the ancestral S-subgenome and 99.7 percent of sequence belong to the ancestral T-subgenome, based on comparison to the subgenome assignments made using the diploid progenitor genomes. The intersection of polyCRACKER classification and classification based on the diploid progenitor genomes constituted 86.5 percent of the total *N. tabacum* assembly, with the remaining sequence almost entirely unclassified by either method (very short scaffolds). We graphically show the high similarity between polyCRACKER and reference-genome based subgenome classification in Fig. 3b,c.

**Figure 3.**
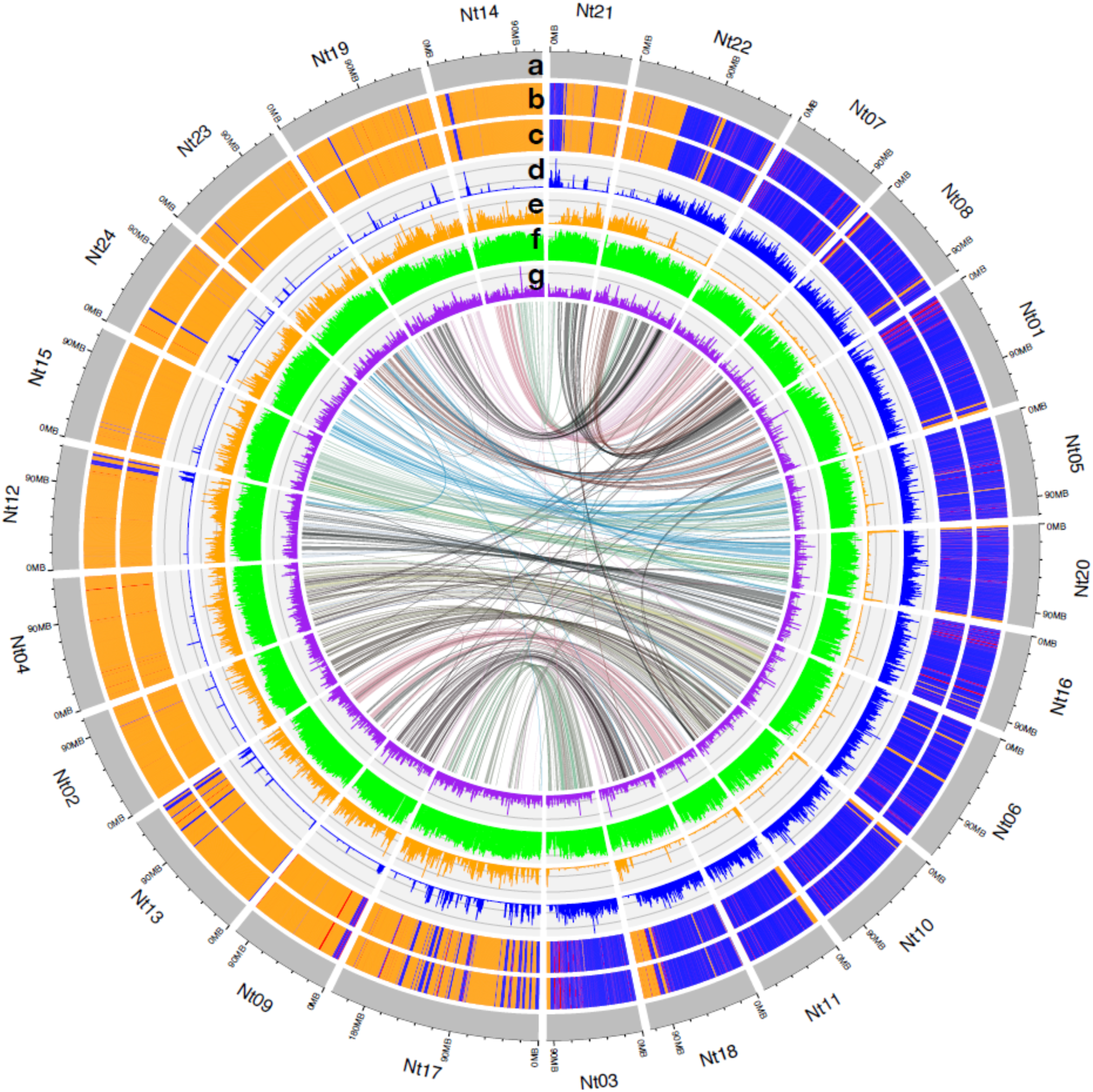
Comparison of *N. tabacum* subgenome classification via polyCRACKER versus progenitor-dependent species binning method. Circos plot showing the 24 pseudomolecules (Nt1–Nt24) in the *N. tabacum* reference genome. With tracks for (**a**) chromosome, (**b**) progenitor-based subgenome classification (**c**) polyCRACKER subgenome classification (d) coverage of polyCRACKER-identified differential k-mers belonging to *N. sylvestris,* (**e**) coverage of *N. tomentosiformis* differential k-mers, (**f**) TE density, and (**g**) gene density. Synteny between pseudomolecules is represented by colored lines within the circle. Tracks **b, c, f,** and **g** represent classification labels and densities spanning 250 kb bins, while tracks **d** and **e** represent coverage over 75 kb bins.

We explored k-mer length as a parameter for polyCRACKER performance and found k-mers as short as 15 nucleotides were sufficient to confidently distinguish the tobacco subgenomes (Fig. 4, Table 1). Increasing k-mer length (specificity) can decrease sensitivity by decreasing the frequency of repeat-k-mers spread across the genome sequences, but this can in part be compensated for reducing threshold for defining a k-mer as repetitive or by reducing the k-mer length (Fig. 4, Table 1). Sensible recommendations for selecting this k-mer length, amongst other hyperparameters, can be ascertained by observing the number of repetitive k-mers per fragment of the fragmented *N. Tabacum* assembly (Fig. 5). If the number of repetitive k-mers is extremely skewed left, as in the two fragmented algae assemblies (Supplemental Figure 1), then the k-mer length should be decreased, at the expense of introducing more noisy degenerate k-mers.

**Figure 4.**
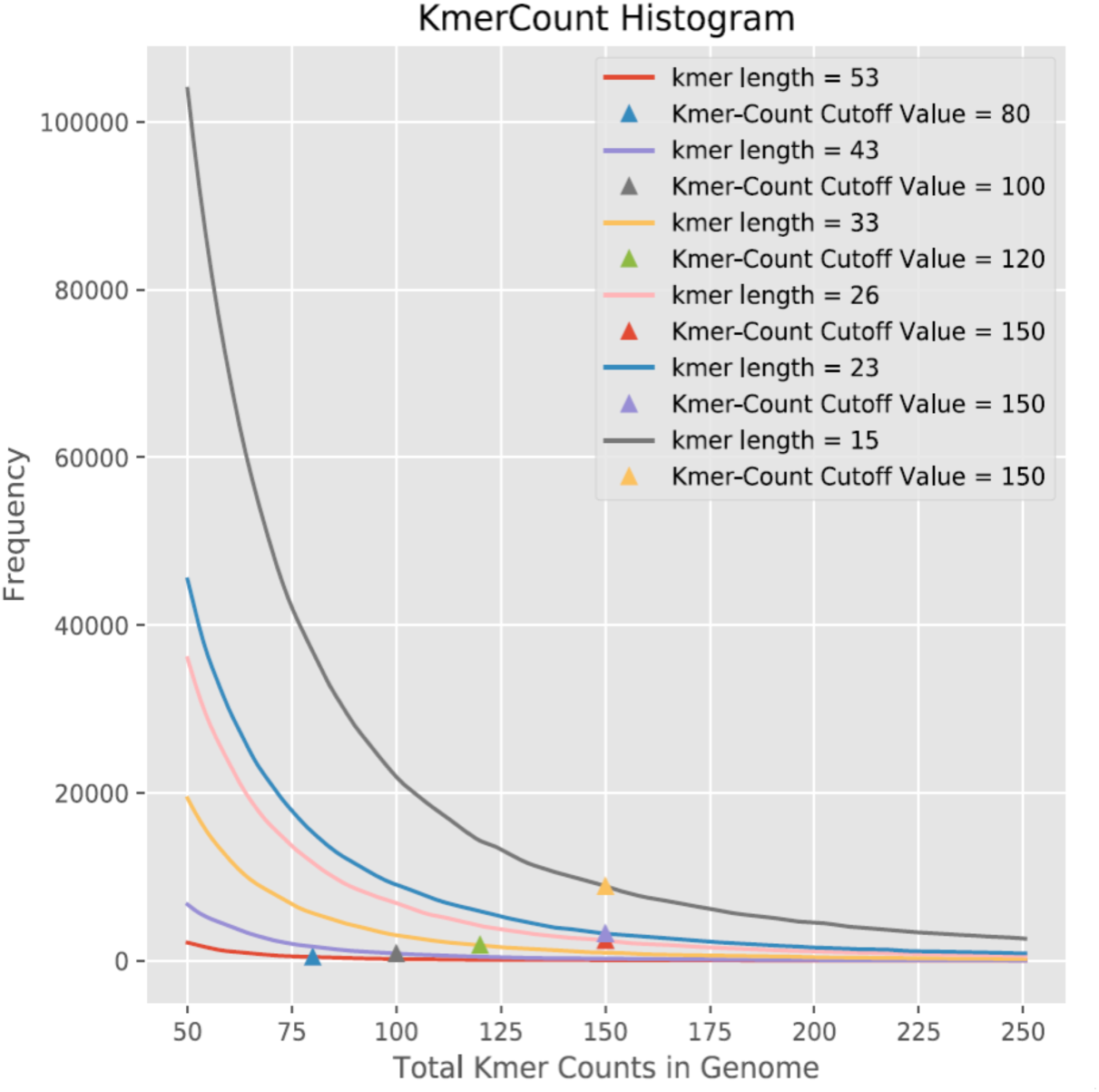
Increasing k-mer length decreases the frequency of repetitive-k-mers spread across the genome. A range of k-mer lengths were tested for *N. tabacum.*

**Figure 5.**
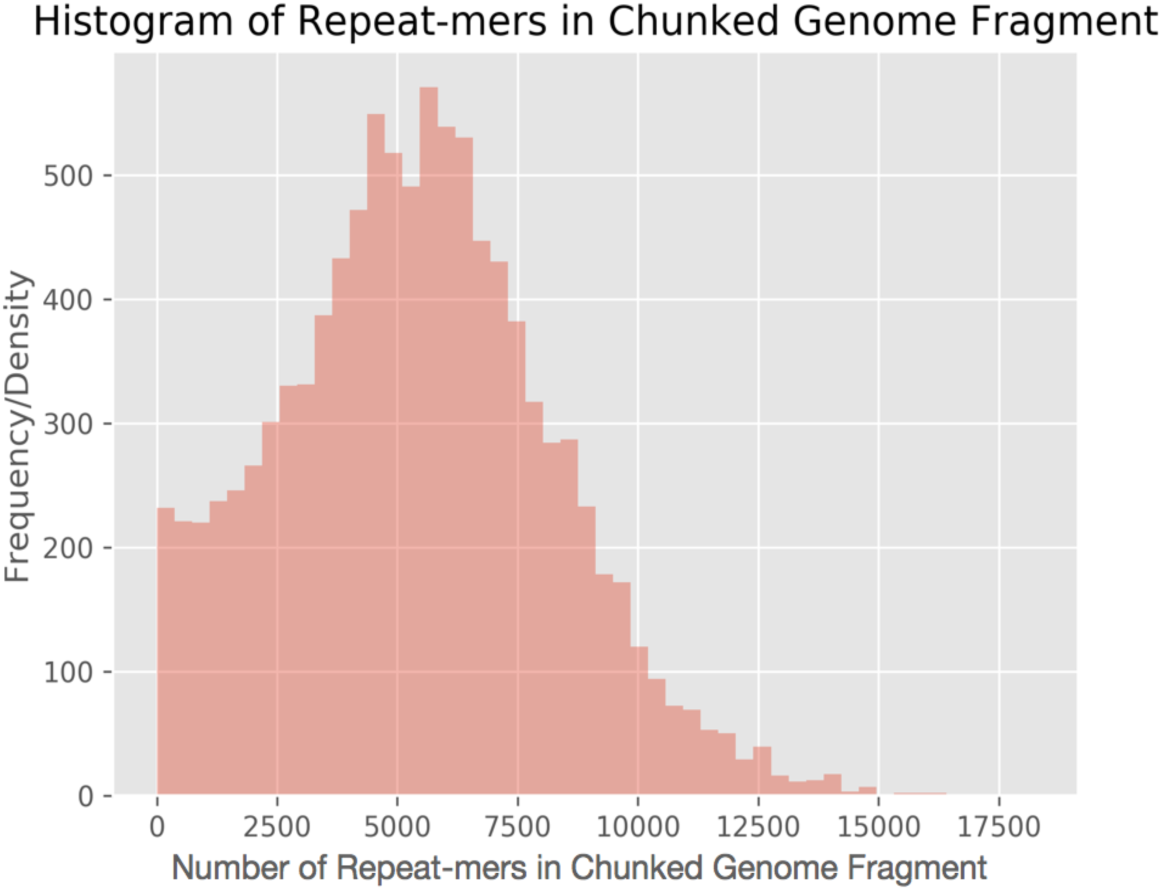
Distribution of the number of repeat-mers for all of the chunked genome fragments of *N. Tabacum*’s psueodomolecule-anchored assembly.

The *N. tabacum* analysis described above was based on the subset of sequences anchored into pseudomolecules [7] in order to show the classification agreement distributed across the chromosomes. However, only 64% of the assembled *N. tabacum* sequence is assigned to pseudomolecules. In order to both provide a comprehensive analysis of *N. tabacum* subgenomes and to show that polyCRACKER work efficiently and accurately on fragmented draft assemblies, we repeated our analysis on the Illumina/454 draft genome assembly from the same study. This draft assembly was produced prior to long-range scaffolding with optical mapping and has a contig L50 of only 8.9 kb (scaffold L50 of 278 kb). Although this assembly was already composed primarily of many small subsequences, we still split larger scaffolds of this assembly into non-overlapping 100 kb segments in order to partially normalize subsequence lengths. polyCRACKER works best with datasets that do not have wide dispersion in length. Subsequences are later easily tracked back to their origin. We then combined subsequences with all scaffolds greater than 2.5 kb. Scaffolds less than 2.5 kb were excluded because they do not contain enough repetitive k-mers for confident classification. After removing the smallest sequences, this dataset contained 94 percent of the *N. tabacum* genome assembly, and PolyCRACKER was able to assign 99.5 percent of sequence estimated to belong to the S subgenome (2.1 Gb, 990 Mb more sequence than that anchored to psuedomolecules) and 99.1 percent of sequence estimated to belong to the T subgenome (1.7 Gb, 262 Mb more sequence than that anchored to psuedomolecules). Sequence assigned to the same subgenome by both polyCRACKER and comparison to the progenitor species constituted 99.2 percent of the genome (Table 2).

### Subgenome classification in the complex allohexaploid, *Triticum aestivum*

To show the scalability of our method, we applied polyCRACKER to group sequences by species of origin for wheat, *Triticum aestivum* [9]. The enormous allohexaploid wheat genome is approximately five times larger than the human genome and its three subgenomes are designated A, B, D. Diploid species similar to the progenitors of two of the subgenomes have been identified: *Aegilops tauschii* for the D subgenome and *Triticum urartu* for the A subgenome. A chromosome-scale genome assembly of *Aegilops tauschii,* the D-genome progenitor of wheat, was recently published [19] and assembled scaffolds are available for *T. urartu* [20]. We used an alignment-based method to assign progenitor of origin to 250 kb non-overlapping segments of the 15 Gb *T. aestivum* assembly [9]. We used this draft genome of *T. aestivum,* rather than the more recent chromosome-scale assembly, as the draft genome is still a relatively fragmented genome (N50 contig size of 232,659 bases), which is reflective of more typical of assemblies for large, complex, allopolyploid genomes. We used 250kb subsequences for initial clustering and included all possible smaller sequences later via signal amplification. PolyCRACKER grouped 12.5 Gb of sequence into one of three sequence bins. Overall agreement between polyCRACKER groups and classification based on alignment to *Aegilops tauschii* and *Triticum urartu* was 0.997 (Cohen’s Kappa) (Table 2).

In Fig. 6c, the PCA plot of scaffolds versus repeat-k-mers (Fig. 6a) were subset and colored by scaffolds belonging to the polyCRACKER-identified A and D genomes. There is a high correspondence between this plot and a PCA plot with the same set of scaffolds colored by their reference-based classification (Fig. 6d).

**Figure 6.**
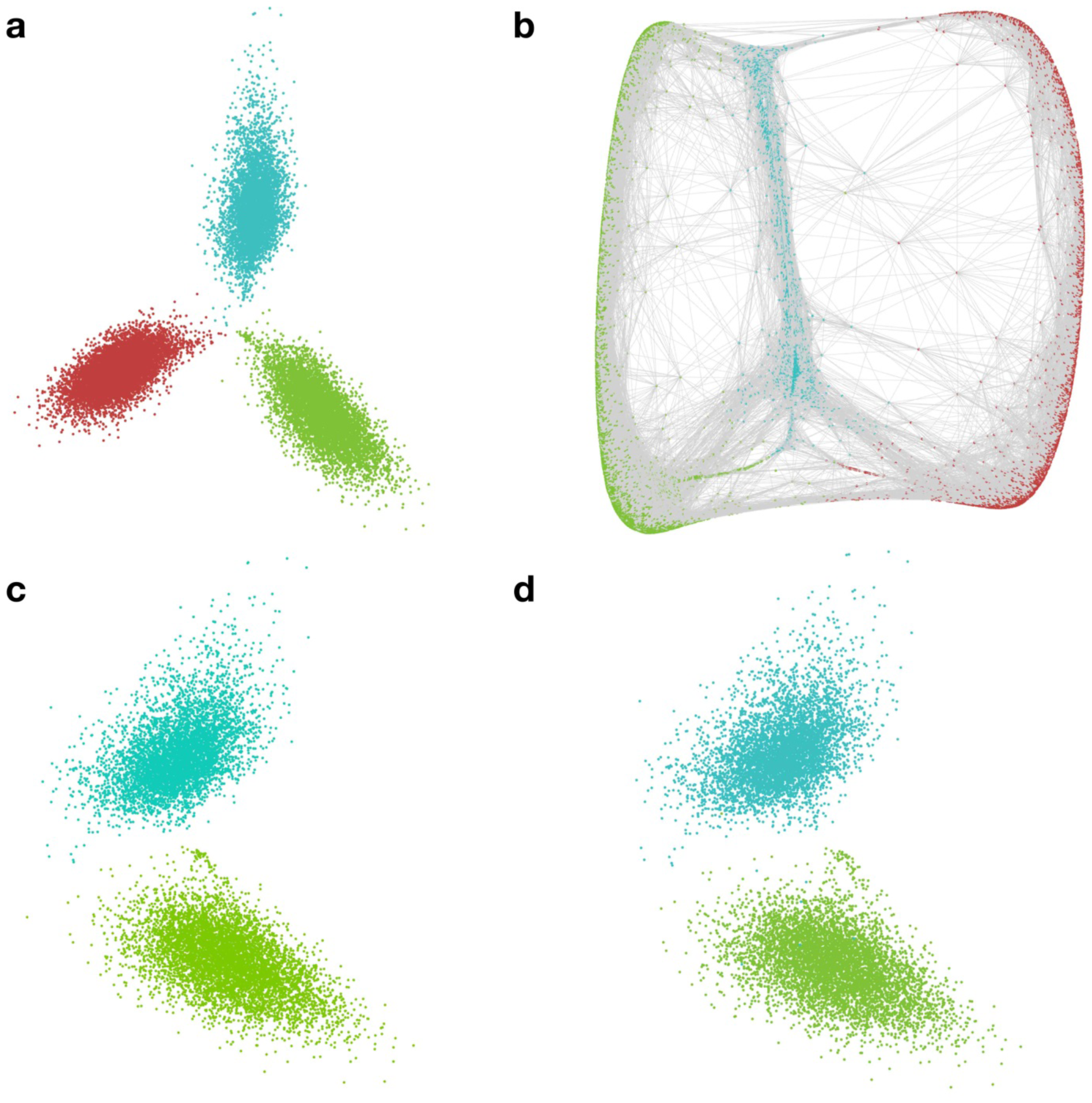
Unsupervised grouping of *Triticum aestivum* subsequences by progenitor of origin using repeat-k-mers. (**a)** PCA labeled by database-free grouping of 250 kb genome segments by polyCRACKER (Subgenomes A, D and B are colored green, blue, and red respectively). (**b)** Spectral embedding of *Triticum aestivum* dimensionality reduced information (reduction to 4-dimensions, visualized in 3-dimensions), in which edges (grey lines) represent shared repeat-mer profiles that connect genome sequences. (**c)** Database-free grouping of A and D subgenomes by polyCRACKER, and (**d**) sequences colored by their similarity to *T. urartu (green)* and *A. tauschii (blue).*

### Repeat annotation and analysis in *N. tabacum*

In addition to using k-mers to separate subgenomes, PolyCRACKER can use repetitive elements, such as LTRs and other transposons, found via common repeat finding programs such as RepeatModeler and RepeatScout, to separate subgenomes by identifying differential repetitive elements.

Using the pseudomolecule-anchored scaffold set, chosen because of a lower dispersion in subsequence length as compared to the unanchored scaffolds, 300 differential repeats were identified, then used to separate the subgenomes of *N. tabacum* following the same approach used for k-mers (Fig. 2d-f). A total of 2.3 Gb out of 2.9 Gb, 79.5% of the total pseudomolecule-anchored genome sequence, was successfully assigned to the S and T subgenomes that were also in agreement to progenitor-mapped labels. The subgenome assignments from the repeat analysis demonstrated 0.87 agreement with the progenitor mapped labels (Cohen’s Kappa) (Fig. 2 d-f).

In addition, polyCRACKER identified enriched subclasses of repeat elements that contributed significantly to the subgenome assignments by comparing the categorical distribution of the informative differential consensus repeats to the null distribution, and assigning a high chi-squared value to repeat subclasses that are over or under represented in the group of informative differential repeats. The top subclass, Unknown, with a highest chi-squared value of 780.74, was found. The same analysis was done for top subclass two, Simple Repeats, and top subclass three, LTR/Gypsy (Supplemental Table 3).

**Table 3.** Dimensionality reduction and clustering techniques used in polyCRACKER for this study.

| Species | Dimensionality reduction technique | Clustering technique |
| --- | --- | --- |
| *N. tabacum* (including unanchored sequences) | KPCA with cosine kernel | Bayesian Gaussian Mixture Models |
| *N. tabacum* (only anchored sequences) | KPCA with cosine kernel | Spectral Clustering |
| *Triticum aestivum* | KPCA with cosine kernel | Spectral Clustering |

## Discussion

The ability to accurately separate closely related genomes or subgenomes without prior knowledge of their composition, relationship or a database of known species is a significant advancement for the study of polyploid genomes. We analyzed test datasets comprised of multiple closely related species and real polyploid genomes to demonstrate the ability of polyCRACKER to separate genomes ranging from small fungal genomes with little repetitive DNA (Supplemental Table 1) to enormous repeat-rich polyploid plant genomes (Table 2). By enabling subgenome-specific analysis polyCRACKER can accelerate rapid advancements in our understanding of polyploid genome evolution and regulation. In particular, it will enable functional studies of genes according to subgenome, promote a deeper understanding of genome dominance and enable comparative studies of diverse polyploids to discern any rules governing polyploid genome evolution.

PolyCRACKER will be particularly useful for the latter by allowing analysis of polyploid species that do not have advanced genetic resources like genetic maps. It should be noted that our overall primary goal for developing polyCRACKER is beyond simply assigning sequences to their subgenome within the modern allopolyploid. Assignment of sequences to their location within the modern allopolyploid can be accomplished by ever-improving technologies including optical maps, genetic maps, Hi-C data, and various long-read sequencing technologies. Rather, the goal of polyCRACKER is to assign sequences to their ancestral progenitor subgenome and clarify what sequences were donated by which ancestral progenitor (even a possibly extinct progenitor). PolyCRACKER is not limited to the analysis of only genomes with long-range contiguity or scaffolding, as demonstrated by our use of fragmented draft genome assemblies. However, if the analyzed genome does have sufficient long-range information, additional insight can be gleaned in the form of identification of translocations between the original progenitor genomes. As shown in Figure 3, modern chromosomes of tobacco are a mosaic of the original progenitor genomes. While this is known for tobacco, due to the availability and prior identification of its progenitor species, for many allopolyploid species the progenitors of origin are unknown or extinct. Therefore, polyCRACKER may be particularly useful in identifying historical or ongoing genetic exchanges between homeologous chromosomes when applied to genomes with long-range information. In such cases, polyCRACKER can uncover the frequency and location of chromosomal translocations between ancestral progenitor genomes. Nonetheless, biological translocations within the evolutionary history of a species is not the only means by which chromosomal mosaics of ancestral progenitors is observed. Indeed, errors in genome assembly of allopolyploids in which a subsequence of one homeologous chromosome is inserted in place of the true sequence often produces the same observation. This occurs frequently in allopolyploid genome assemblies as many sequences within homeologous chromosomes are virtually identical (with the exception of transponsons, viral DNA and other repetitive sequences). Thus polyCRACKER may be useful check for evaluating assembly accuracy and identifying potential mis-assemblies in allopolyploid genomes.

Repetitive DNA is a major component of eukaryotic genomes. While polyCRACKER exploits the rapid turnover and evolution of repetitive elements to separate closely related genomes, it can also be used as tool to study the origin, evolution, and impact of repetitive DNA on the entire genome. We used the differential k-mers identified by polyCRACKER to assign assembled repeats to subgenomes. This also placed the k-mers into the context of intact repetitive elements. We also demonstrated that subgenome-specific repetitive elements could be used to bin subsequences by subgenomes.

Existing unsupervised methods used to separate subgenomes were developed to bin genomes from metagenome samples and are ill-suited for classifying allopolyploids because the homeologous chromosomes often have more similar overall k-mer profiles than the profiles of chromosomes within their respective subgenomes. In contrast to previous algorithms and approaches for grouping sequences by species of origin, we focus on the fact that even closely related species often differ in the frequency of repeat sequences present throughout their genome. This is particularly useful for polyploid genomes because such genomes are usually so closely related that they do not differ in coarse genome features like tetranucleotide frequency, GC content, or codon usage. Such repeats may correspond to sequences associated with transposons, viruses, and other parasitic sequences that have proliferated in one species relative to another. By 1) counting the number of instances of short and/or long repeat k-mer sequences, 2) creating a sparse matrix of number of instances of k-mers by the subsequences on which they occur, 3) reducing the dimensionality of the sparse matrix by one of several known (Table 3), and 4) applying a clustering method (Table 3) we can group scaffolds into bins that correspond to distinct species. We can then further increase signal by taking the resulting binned sequences and 1) counting k-mer frequency, 2) identifying k-mers that are enriched in one bin versus other bins, and 3) use the presence of enriched k-mers on previously ambiguous sequences to group them into the appropriate species bin.

Biased loss of repetitive sequence is often observed in allopolyploids and in the case of *N. tabacum,* repetitive sequence appears to have been preferentially lost from the T-subgenome [17]. Indeed, both polyCRACKER and database-dependent classification of *N. tabacum* confirm that the S-subgenome is substantially larger than the T-subgenome as suggested by others previously, and consistent with biased repetitive sequence loss from the T subgenome. We also identify several classes of repeats that are enriched in each subgenome. Our observation of significant enrichment of simple repeats in the T-subgenome is consistent with the previous observation that some classes of satellite repeats are several-fold more prevalent in *N. tomentosiformis* than in *N. sylvestris* [17, 21]. Our observation of significant enrichment of tranLTR/Gypsy and unknown elements in the S-subgenome of *N. tabacum* is consistent with the prior suggestion that *N. sylvestris,* but not *N. tomentosiformis,* had recent bursts of repeat element proliferation, likely involving Gypsy transposable elements [17]. In conclusion, polyCRACKER is a robust method for classifying sequences according to species of origin that will be important in future studies of allopolyploids.

## Methods

We describe a method to group sequences belonging to the same subgenome that consists of connecting sequences within a graph in which edges are shared profiles of repetitive k-mers. The network graph is established by considering a user-supplied number of regions with closest repeat distributions (number of nearest neighbors). The distance between sequence nodes in the graph is indicative of the level of shared k-mers. Thus, polyCRACKER groups sequences by species of origin by connecting nodes, representing sequences, by edges that represent the common occurrence of repetitive DNA sequences unique to that species or subgenome. This technique was applied to highly fragmented draft genome assemblies or simulated assemblies. Scaffold lengths were normalized by splitting larger sequences into shorter subsequences. PolyCRACKER identifies potentially informative k-mers that occur above a minimum frequency. This greatly reduces the number of k-mers that are subsequently mapped to the scaffolds to determine their per scaffold frequency.

PolyCRACKER creates a sparse matrix, in which rows correspond to fragments and columns are unique k-mers, and each intersection contains the frequency of each unique k-mer on each scaffold subsequence. Then, it performs dimensionality reduction on the sparse matrix, projecting the high-dimensional data into a lower-dimensional space, where each subsequence is represented by a point in the lower dimensional space. At this point we can visualize the data in three-dimensional graphical plots (Figs. 2c,f and 6b). In order to assign sequences to species or subgenomes, polyCRACKER performs unsupervised clustering on the low-dimensional data via clustering. Any ambiguous sequences are removed, but may be used later through a signal amplification method described below.

To assign additional subsequences to species bins, polyCRACKER identifies highly differential k-mers between the initially grouped scaffolds and uses these k-mers to recruit the remaining, ambiguous subsequences. Similar to above, we identify unique k-mers for each preliminary group of scaffolds, and then identify k-mers that differentiate those already grouped sequences by comparing the number of occurrences of a particular k-mer in one group versus the others. K-mers that occur frequently in one group, but not in at least one of the others, past a certain threshold, are output to a FASTA file as differential k-mers for that group. PolyCRACKER maps each set of the differential k-mers against the entire set of scaffolds in the assembly. We then find the total counts for the sum of all differential k-mers of a particular group for each subsequence. The results of the aforementioned step may be plotted across entire chromosomes (Fig. 3) as a cross-check for the test case, which was done using shinyCircos [22] (polyCRACKER formats the input data for shinyCircos). When the total counts of an inferred subgenome’s differential k-mers in a region are substantially larger than that of the differential k-mers of another inferred subgenome in that region, polyCRACKER extracts the binned sequences and stores them in FASTA format. The now larger set of grouped subsequences can be used as the new input for another round of differential k-mer analysis to recruit more subsequences. The previous steps (recruiting differential k-mers and binning sequences) are repeated until the number of iterations reaches a user-defined cutoff, and the user can select the species bins from any iteration of the binning process. The final results and spectral graph can be visualized in several ways, including force-directed physics simulations of the *K-nearest-neighbors* graph to visualize changes in the data’s structure over time. An analogous approach can be taken for binning sequences by repeat elements, although the k-mer analysis results must be used as an input for that algorithm to identify the initial differential repeat elements.

Itemized below is a high-level flow summary for the aforementioned methodology:

1. Break the assembly into chunks of a fixed size, keeping track of the scaffolds from which these chunks came from.
2. Bin the chunks by subgenome.
3. Optionally, merge the binned chunks back together based on their scaffold of origin (or bin the entire original scaffold based on the total counts of differential repetitive sequence). In addition, there is an option available to assign remaining unassigned fragments/scaffolds to a subgenome bin based on the consensus assignment of all its neighboring/associated fragments.

### Recommendations for Selection of PolyCRACKER Hyperparameters

While the search space of polyCRACKER’s hyperparameters remains fairly unexplored, the algorithm provides a few functions that can make recommendations on how to set the hyperparameters to achieve reasonable performance. Of vital importance is making sure that the parameters do not result in too few or too many repeat-kmers in the analysis. If there are too few repeat-kmers included in the genome fragments, then they will be hard to bin. The function *number_repeatmers_per_subsequence* plots a histogram of the number of repeat-mers present in each chunked genome fragment. If this histogram is skewed to lower k-mer counts in each fragment, then reducing the k-mer size, increasing the length of the chunks, decreasing the minimum genome-wide k-mer frequency for k-mer inclusion, amongst other changes can help increase the repetitive content of each fragment, thereby improving the initial binning of these fragments into subgenomes. There exists help documentation in the polyCRACKER source code repository that can help inform about sensible binning practices.

### Propagating PolyCRACKER Labels via the Relative Position of Subsequences

Since polyCRACKER performs best when the length of the input scaffolds are similar, we split scaffolds into fragments of similar length as we did when we prepared the *N. tabacum* input dataset. However, it is still possible to use prior information about the location of the fragments in scaffolds. For example, if one or several fragments belonging to a scaffold are classified to a subgenome, it is likely that other subsequences from the same scaffold are also associated with that same subgenome. PolyCRACKER uses this information by passing subgenome labels of classified subsequences through a semi-supervised label propagation algorithm that takes into account labels as well as relative genomic positions from which subsequences are derived to infer the labels of the unclassified subsequences. Conversely, when interrogating unvalidated assemblies, discordance between scaffold labels and species assignment may indicate assembly errors that can be fixed by breaking scaffolds.

### Random sampling of repeat k-mers for datasets too large to include all k-mers

PolyCRACKER can randomly subsample repeat k-mers when the data set is too large to include all k-mers with available computational resources. For example, there were 550,790,662 unique k-mers with frequencies between 3 and 100 within the *Triticum aestivum* genome. A matrix 550,790,662 columns by many thousand rows is computationally intensive to construct and subsequently analyze. We therefore randomly sampled 2,420,450 k-mers to generate initial partitions of the subgenomes, and further subsampled differential k-mers to obtain 568,637 k-mers for the final partitioning of *Triticum aestivum* subgenomes.

### Subgenome validation for complex allopolyploid species

For validating polyCRACKER subgenome groups in the context of the *N. tabacum* and *Triticum aestivum* genomes in which subgenomes were not already labeled, we used the progenitor species to assign species of origin to respective sequences. The PolyCRACKER *progenitorMapping* function exploits *seal.sh* function within the bbtools toolkit (sourceforge.net/projects/bbmap/) to bin subsequences based on reference progenitors, and serves as a tool to compare polyCRACKER to reference-based binning solutions. Progenitor mapped labels and binned sequences are output from this analysis and can be visualized on the PCA or the spectral embedded data. If polyCRACKER is used to separate species from an input genome instead of subgenomes, the original species labels for the subsequences can be recovered.

### Quantitative analyses on the classification and clustering polyCRACKER results

The polyCRACKER function *final_stats* was used to perform quantitative analyses on polyCRACKER’s classification and clustering results, with comparisons to the results achieved by the reference-based *progenitorMapping* and to the ground truth species extraction with known species. For a classifier comparison of polyCRACKER’s results to the progenitor mapped subgenomes, the lengths of each subgenomes from both analyses were found and compared to each other and the original progenitor subgenome sizes, and amongst other measures, Cohen’s Kappa and Jaccard Similarity between the amount of unambiguous sequence shared between both analyses were calculated as measures of agreement between the reference based and unsupervised methods. For accuracy measures of polyCRACKER versus ground truth labeling, such as when the subgenomes or species of each region/chunk are already known, the amount of sequence correctly classified, precision, recall, and f1 scores were reported for each species/subgenome and averaged, amongst others. Confusion matrices for each of the two classification test types are output from the analysis. Analyzing the clustering results, total sequence lengths for each cluster is found and Silhouette and Calinski Harabaz scores can be calculated if supplying the original PCA data along with the final polyCRACKER output labels. If either ground truth labels or progenitor mapped labels are supplied, some measure of agreement, homogeneity, and completeness between the two results can be found.

### Repeat annotation and analysis in *N. tabacum*

PolyCRACKER studies annotated repeats that are identified using RepeatModeler, which uses the de-novo repeat finding programs RECON [23] and RepeatScout [24] to employ complementary computational methods for identifying repeat element boundaries and family relationships. RepeatModeler builds, refines and classifies consensus models of putative interspersed repeats from RECON and RepeatScout output. We further filtered repeat annotations using Pfam, and PANTHER annotations of predicted repeats. The final non-redundant de novo repeat database was then used with RepeatMasker to annotate repeats across the genome sequence.

### Subgenome classification using genome-wide annotated repeats in *N. tabacum*

Genome-wide RepeatMasker annotation of repeats was used with *TE_cluster_analysis* to generate a sparse matrix of the subsequences by their distribution of repeats belonging to a particular family, family member, class, and subclass (had a specific consensus sequence label) via *repeatGFF2Bed* and *sam2diffk-mer_clusteringmatrix.* Each row of this matrix was labeled by polyCRACKER’s k-mer analysis final subgenome labeling, and the highly informative repeats were found by calculating the chi-squared statistic of each column of the matrix. PolyCRACKER transformed this matrix into a matrix depicting the total counts of a particular repeat for each subgenome (the differential repeat subgenome matrix), and labeled a repeat by a subgenome if it was differentially present in one subgenome above a user-specified threshold. The consensus repeats were assigned a higher chi-square statistic if their assembly frequency was indicative of greater dependence between repeat and species label and polyCRACKER denotes these repeats as highly informative. The consensus repeats with the highest chi-square statistic were chosen from the differential repeats to find the most informative differential repeats, and the same number of repeats were selected from each subgenome for subsequent analysis. To cluster and extract the subgenomes using a repeat count matrix, this matrix was fed into *subgenome_extraction_via_repeats,* which finds the most highly informative differential repeats, feature selects the matrix by these repeats, performs dimensionality reduction using PCA, finds a neighborhood community graph and clusters the resulting dataset via Spectral Clustering. Signal amplification is employed on these clusters analogous to the k-mer analysis in order to iteratively recruit more subsequences to the final bins. PolyCRACKER also employs multiple algorithms to discover repeat element subclasses that are important and informative for differentiating the subgenomes, as discussed in the Supplementals.

### Distinguishing species of origin using *de novo* repeat annotations

PolyCRACKER grouped *N. Tabacum*’s pseudomolecule-anchored assembly (broken into 250 kb fragments) by species of origin using *de novo* repeat annotations in a similar method as used for k-mers. A matrix of scaffolds vs repeats was constructed. Differential repeats were identified as defined by a five-fold greater presence in one subgenome versus the other. Subgenome labels found via polyCRACKER’s k-mer based binning were then used to identify highly informative repeats (features with high chi-squared value). Intersection of differential repeats with the 150 most informative repeats from each subgenome yielded 300 informative differential repeats. PolyCRACKER then performed feature selection on the matrix, keeping columns/repeats corresponding to highly informative repeats. PolyCRACKER performed principal component analysis and clustered the feature selected matrix to find new labels of scaffolds and to see if repeats can distinguish subgenomes. In this clustering process, nodes, representing scaffolds, were linked by edges that connected two nodes with similar consensus repeat coverage/content, and Spectral Clustering was used to perform normalized cuts of this graph to form initial partitions. Using the same process as outlined for k-mers, differential repeats were identified between the initial sequence bins by being eight or greater times present in one genome versus the other, and then they were used to recruit more subsequences via the signal amplification process, finding the total hits of these differential repeats in each bin. PolyCRACKER reassigned subgenome labels if their ratio of hits of the total differential repeat counts is greater than a threshold, in this case 3 times greater. The process of identifying differential repeats and then re-binning by the ratio of total repeat counts for each scaffold is reiterated via a bootstrap process until convergence, in which after a number of trials, the amount of sequence assigned to each bin does not appear to make large changes, converges on a set amount of sequence, and the number of times this process can be run is left to the user’s discretion.

### Identification of informative repeat classes

Consensus repeats, identified during *de novo* repeat finding and labelled by unique identifiers were identified by polyCRACKER as differential if the number of identified instances of that consensus repeat were five or more times more frequent in one subgenome versus the other. Out of all differential repeats identified, polyCRACKER selected 400 (200 from each subgenome) of these repeats (from the unanchored *N. Tabacum* scaffolds) as subgenome signatures. The repeats were grouped together by their subclass annotation. Specific subclasses of repeats were found by polyCRACKER to be highly informative if their number of informative differential consensus repeats was statistically significantly under or over represented versus a similar distribution of the consensus repeats across the entire genome (via a chi-squared statistic).

## Declarations

### Availability of data and material

The polyCRACKER algorithm is available at: https://bitbucket.org/pangenome2/polycracker/

## Competing interests

The authors declare no competing interests

## Funding

The work conducted by the US DOE Joint Genome Institute is supported by the Office of Science of the US Department of Energy under Contract no. DE-AC02-05CH11231. This work was supported by the Laboratory Directed Research and Development Program of Lawrence Berkeley National Laboratory under U.S. Department of Energy Contract No. DE-AC02-05CH11231

### Authors’ contributions

SPG conceived of the implementation and overall study design, wrote a prototype algorithm to use known differential repeat-k-mers to recruit sequences into respective species/subgenome bins, developed some initial visualization methods, and performed bioinformatic analyses during software development and for identification of consensus repeats; JL introduced the unsupervised binning approach to the study design, improved the supervised approach, conceived and implemented the use of sparse matrices, dimensionality reduction and clustering techniques, designed the figures and methods, wrote the program, designed and optimized the bioinformatics workflows and algorithms for high-performance computing and developed data visualization methods, and performed the analyses for the aforementioned assemblies; JPV provided overall project supervision and helped interpret results. All authors contributed to writing the manuscript.

## Supplementary Material

### polyCRACKER performance on simulated datasets comprised of mixtures of subsequences from multiple closely-related species

Three simulated datasets created from reference fungal or algal genomes were used to develop and test polyCRACKER. The genomes were chosen to cover a range of repeat contents and genomes sizes. The simulated datasets were created by fragmenting and pooling the reference genomes to simulate the contigs expected from a very simple metagenome or polyploid genome. PolyCRACKER was then optimized to correctly assign the sequence fragments to their genome of origin.

The first simulated dataset contained genomes from two closely related (mean nucleotide identity of CDS, 77%) Basidiomycte fungi, *Ustilago hordei* [17] and *U. maydis,* whose small (~20Mb) genomes contain only 7.8% and 2% repetitive DNA, respectively [17, 18]. Genomes of the two species were broken into 50 kb fragments, labeled by species (for subsequent calculation of accuracy), and then combined into a single FASTA file. Without any other input, polyCRACKER assigned sequences to the correct species with perfect precision and 99% recall (Table 1).

**Supplemental Table 1.**
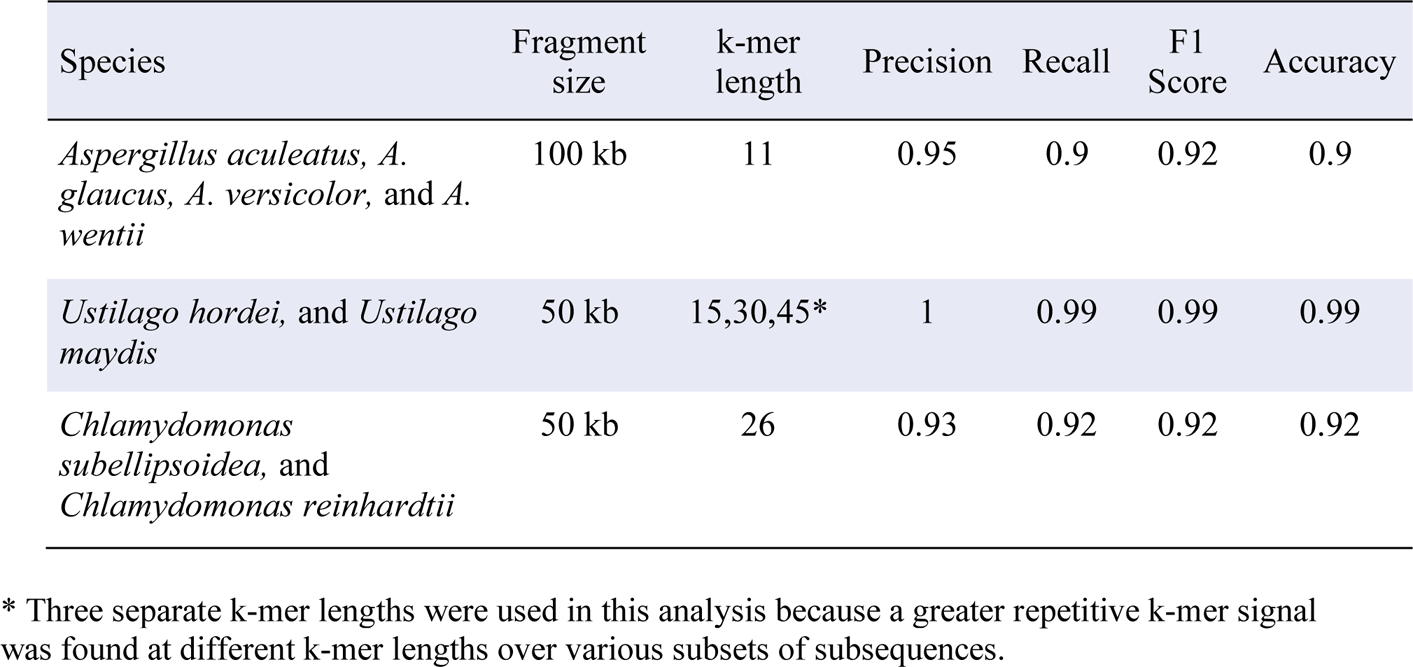
Unsupervised separation of simple mixtures of fungal and algal genomes using PolyCRACKER’s differential k-mer analysis.

| Species | Fragment size | k-mer length | Precision | Recall | F1 Score | Accuracy |
| --- | --- | --- | --- | --- | --- | --- |
| *Aspergillus aculeatus*, *A. glaucus*, *A. versicolor*, and *A. wentii* | 100 kb | 11 | 0.95 | 0.9 | 0.92 | 0.9 |
| *Ustilago hordei*, and *Ustilago maydis* | 50 kb | 15,30,45* | 1 | 0.99 | 0.99 | 0.99 |
| *Chlamydomonas subellipsoidea*, and *Chlamydomonas reinhardtii* | 50 kb | 26 | 0.93 | 0.92 | 0.92 | 0.92 |

\* Three separate k-mer lengths were used in this analysis because a greater repetitive k-mer signal was found at different k-mer lengths over various subsets of subsequences.

**Supplemental Table 2.**
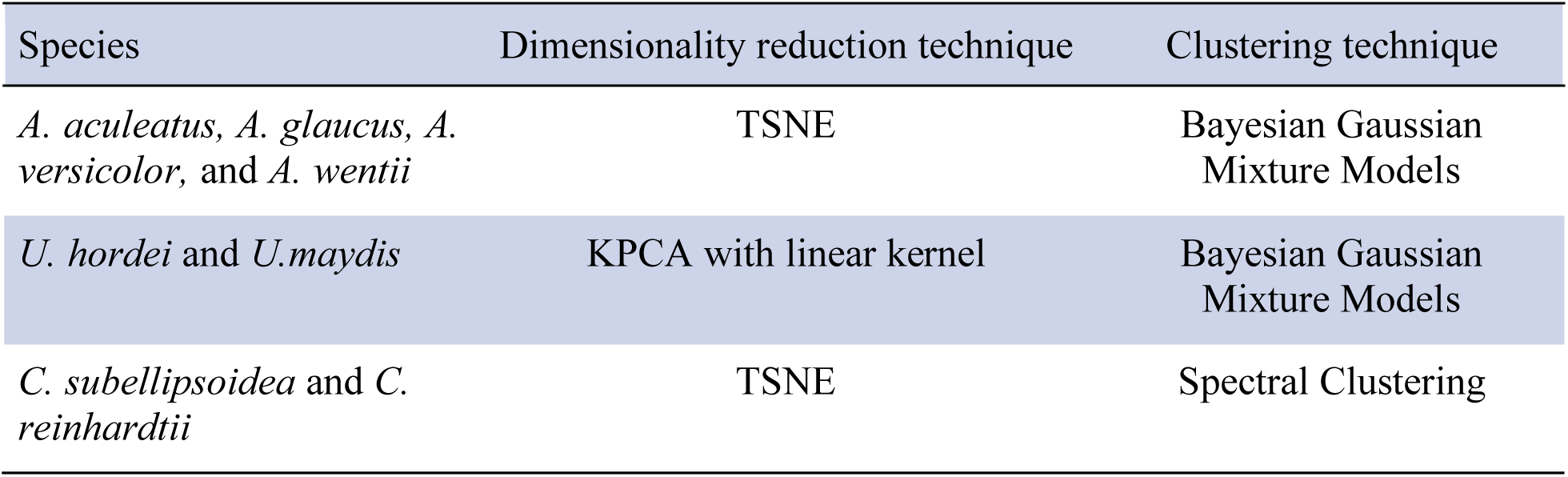
Dimensionality reduction and clustering techniques used in polyCRACKER for the simulated metagenomes.

The second simulated dataset contained four closely related (pairwise nucleotide identity of CDS ranged from 63-70%) Ascomycete fungi: *Aspergillus glacus* (30 Mb, 3% repetitive), *A. aculeatus* (35 Mb, 4.5% repetitive), *A. versicolor* (33 Mb, 1.8% repetitive), and *A. wentii* (34 Mb, 1.8% repetitive) [19]. The four genomes were broken into 100 kb fragments and pooled as for the first dataset. PolyCRACKER assigned the sequences to the correct species with 95 percent precision and 90 percent recall (Table 1).

Lastly, we tested our method on a simulated dataset containing two single-cell green algae: *Chlamydomonas reinhardtii* [20] (16% repetitive DNA) and *Coccomyxa subellipsoidea*] (1.5% repetitive DNA). The simulated dataset was created exactly as for the first dataset. PolyCRACKER assigned the sequences to the correct species with 93 percent precision and 92 percent recall (Table 1).

The dimensionality reduction and clustering techniques used to establish the initial species bins are depicted in Table 2. The amount of repetitive k-mers in each of the genome fragments of the two algae lines’ broken assemblies is shown in Figure 1. It is fairly low as compared to *N. Tabacum’s* distribution of its repetitive content, after scaling for fragment length.

**Supplemental Fig. 1.**
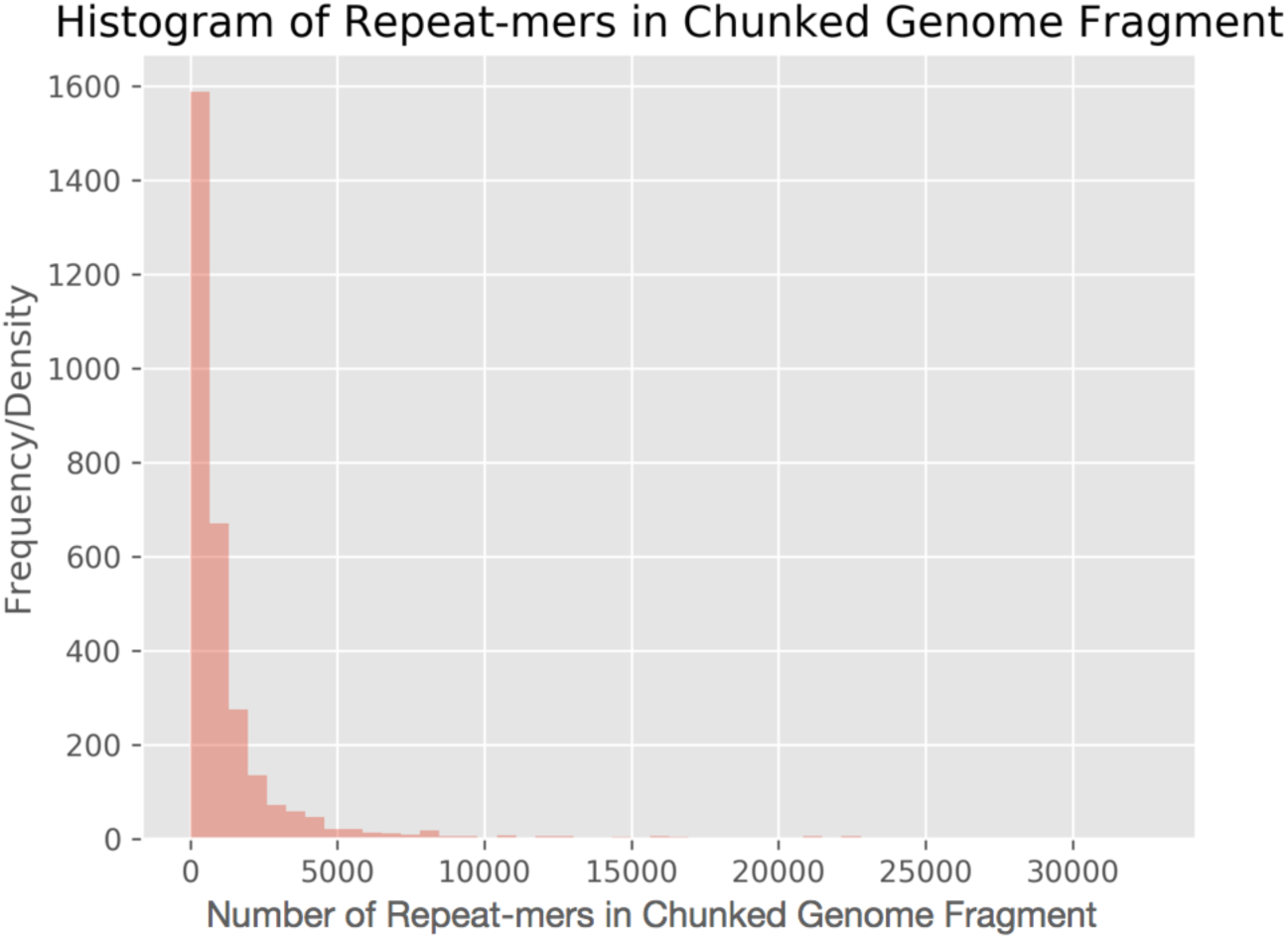
Distribution of the number of repeat-mers for all of the chunked genome fragments of the two single-cell green algae assemblies. These numbers were derived from counting the total number of repetitive k-mers in each genome fragment after breaking the assembly into chunks of a fixed size.

### Highly differential repeat subclasses between *N. Tabacum* subgenomes

Two hundred informative differential repeat consensus sequences were chosen for each subgenome (S and T) of the unanchored scaffold set via the same process used to identify highly informative differential repeats, and analyzed for their subclasses and the subgenome specificity of these subclasses. All repeats and the top differential repeats were broken down into their class or subclass and the categorical distribution of all consensus repeats were used as the null distribution. The categorical distribution of the top differential repeats were compared to the null distribution, and expected number of top differential repeats were found for each subclass. The resulting distributions with the top four chi-squared values are displayed in Table 3. The top subclass, Unknown, with a highest chi-squared value of 780.74, was found, and all top differential Unknown consensus sequences were broken down into their subgenomes and a phylogenetic tree of 25 bootstraps was found between the consensus sequences. The same analysis was done for top subclass three, LTR/Gypsy, but not subclass two, since Simple Repeats are very small and aligning them yields little to no new information about their clade structure. However, it should be noted that Simple Repeats were entirely T subgenome enriched. Information on the subgenome specificity of each top differential subclass and the phylogenetic tree of those sequences can be found in Figure 2.

---

**Supplemental Table 3.** *N. Tabacum* subgenome repeat analysis.

|  | LTR<br>Copia | LTR<br>Gypsy | Simple<br>repeat | Unknown<br>repeat |
| --- | --- | --- | --- | --- |
| Number of consensus sequences | 151 | 527 | 10,694 | 1,414 |
| Number included in the 400 most<br>informative repeats | 5 | 70 | 90 | 229 |
| Expected number among 400 most<br>informative repeats | 4.68 | 16.35 | 331.93 | 43.88 |
| Chi-Squared Value | 0.02 | 175.9 | 176.3 | 780.7 |
| Top 400: Number of S subgenome to T<br>subgenome consensus repeats | 1:4 | 43:27 | 0:90 | 152:77 |

**Supplemental Fig. 2.**
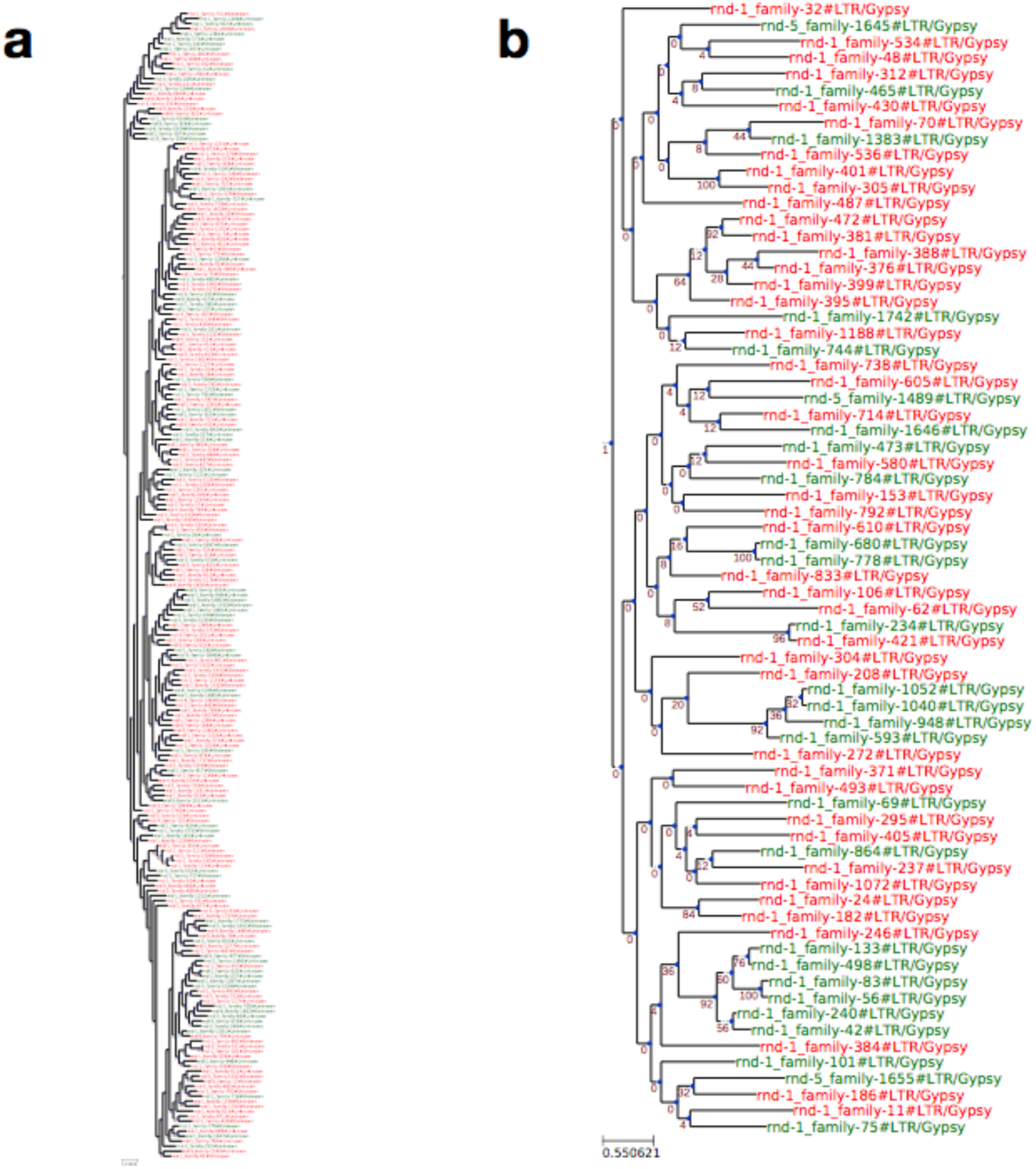
Specific repeat classes contribute to the differentiation of the *N. tabacum* subgenomes. IQTree GTR trees for top subclasses of *N. tabacum* genome assembly. The top three differential classes are **(a)** transposons with unknown class enriched in the S subgenome (red) **(b)** tranLTR/Gypsy (S subgenome enriched). Phylogenetic trees were constructed using maximum likelihood General Time Reversible (GTR) model of evolution, performed with 25 bootstrap iterations.

